# Variant-specific priors in colocalisation analysis

**DOI:** 10.1101/2024.08.21.608957

**Authors:** Jeffrey M. Pullin, Chris Wallace

## Abstract

Linking GWAS variants to their causal gene remains an ongoing challenge. A widely used method for performing this analysis is the coloc package for statistical colocalisation analysis, which can be used to link GWAS and eQTL associations. Currently, coloc assumes that all variants in a region are equally likely to be causal, despite the success of fine-mapping methods that use additional information to adjust their prior probabilities. In this paper we propose and implement an approach for specifying variant-specific prior probabilities in the coloc method. We describe and compare six source of information for specifying prior probabilities: non-coding constraint, enhancer-gene link scores, the output of the PolyFun method and three estimates of eQTL–TSS distance densities. Using simulations and analysis of ground-truth pQTL–eQTL colocalisations we show that variant-specific priors, particularly the eQTL–TSS distance density priors, can improve colocalisation performance. Furthermore, across GWAS– eQTL colocalisations variant-specific priors changed colocalisation significance in up to 13.5% of colocalisations, at some loci revealing the likely causal gene.

## Introduction

Genome-wide association studies (GWAS) have uncovered hundreds of thousands of disease associated variants [1, 2]. However, more than 90% of these variants lie in non-coding regions of the genome, making elucidating their functional consequence challenging [3, 4]. Instead, GWAS associations generally lie in open chromatin regions and are enriched in gene regulatory regions, suggesting they act by modulating gene expression [4, 5, 6]. Therefore, a widely used approach to interrogate variant function is assessing whether GWAS and variants associated with gene expression – expression quantitative trait loci (eQTLs) – are caused by the same variants using statistical colocalisation methods [7]. However, most GWAS hits do not have evidence of colocalisation, representing a large ‘colocalisation gap’ [8].

Today, large amounts of functional annotations and other additional information about genetic variants are available, with initiatives such as The Encyclopedia of DNA Elements (ENCODE) providing genome-wide catalogues of functional annotations. It is well established that GWAS associations are enriched among annotations such as DNase I Hypersensitive sites, implying that functional annotations can provide information about whether variants are causal [4]. More recently, studies have created genome-wide enhancer-gene maps using the activity-by-contact (ABC) score with the intention of prioritising variants that lie in enhancers [10]. Estimates of genomic constraint in non-coding regions have also been generated from large whole genome sequencing datasets, with the idea that highly constrained regions are more likely to have functional consequences [11]. Finally, there are large amounts of publicly available eQTL data that can be used to estimate the most likely positions of eQTLs relative to genes.

Computational methods across statistical genetics have been designed to incorporate this additional variant-specific information to improve performance. For example, in statistical fine-mapping, methods such as PAINTOR [12] incorporate functional annotations in their estimation procedures, while the PolyFun method estimates the prior probability of causality for variants, for use in fine-mapping, based on functional data [13]. A key motivation for using additional data in fine-mapping is that it can ‘break ties’ between variants in strong linkage disequilibrium, which are otherwise impossible to distinguish. Incorporating functional annotation can greatly concentrates the posterior probabilities on a smaller number of variants. In an analysis of 49 UK Biobank traits, use of PolyFun prior probabilities with the SuSiE fine-mapping method [14], lead to 32% more variant–trait pairs with posterior causal probability > 0.95 [13]. Beyond fine-mapping, locus-to-gene models, such as OpenTarget’s L2G model, [15] incorporate a range of genetic information, including distance and predicted variant effects. Incorporating distance information was also recently shown to greatly improve peak-to-gene link prediction [16]. Despite these successes however, to date no methods exists for incorporating additional variant-specific information into statistical colocalisation analyses.

A widely used method for performing colocalisation analysis is the coloc R package, which uses a Bayesian approach to calculate the probability of hypotheses corresponding to different scenarios of shared and distinct causation of associations [7]. Specifically, coloc computes the posterior probability of five hypotheses:

- *H*_0_: no association with either trait;
- *H*_1_: association with trait 1 only;
- *H*_2_: association with trait 2 only;
- *H*_3_: association with both traits, distinct casual variants; and,
- *H*_4_: association with both traits, shared causal variants.

The probabilities are computed by enumerating all possible configurations of causal variants, assuming there is at most one causal variant for each trait. Each configuration has a prior probability calculated from three variant level prior probabilities:

- *p*_1_: the probability each variant is causal for trait 1 only;
- *p*_2_: the probability each variant is causal for trait 2 only; and
- *p*_12_: the probability each variant is causal for both traits.

Previous work has found that the values *p*_1_ = *p*_2_ = 10^−4^, *p*_12_ = 5 × 10^−6^ lead to robust inference over a range of scenarios [17]. In addition, coloc was recently extended using the SuSiE fine-mapping algorithm [14] to relax the assumption of a single causal variant in each region [18].

Currently, coloc assumes that the prior probabilities *p*_1_, *p*_2_ and *p*_12_ are uniform across variants in the region of interest. In this paper, inspired by the success of the use of additional variant-specific information in fine-mapping, and with the ultimate goal of narrowing the colocalisation gap, we propose an implementation of variant-specific prior probabilities in coloc. We describe and compare the effectiveness of various sources of information for specifying these prior probabilities using simulated data and pQTL–eQTL colocalisations. Finally, we demonstrate the effect of using variant-specific prior probabilities on eQTL–GWAS colocalisation analyses.

## Results

### Incorporating variant-specific prior probabilities in coloc

The `coloc.abf(), coloc.susie()` and `coloc.bf_bf()` functions in coloc have been updated to include two new arguments: `prior_weights1` and `prior_weights2`, which specify non-negative weights for the probability of a variant being causal for trait 1 and trait 2 respectively. These weights are used to calculated variant-specific prior probabilities, according to the formula

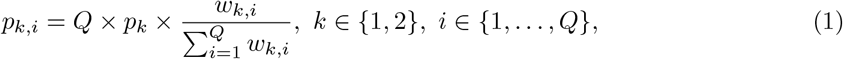

where *w*_*k,i*_ is the provided weight for SNP *i* and trait *k, Q* is the number of variants analysed, and *p*_*k,i*_ is the prior probability of SNP *i* for trait *k*. The variant-specific prior probabilities of being causal for each trait are then used to calculate the variant-specific prior probability of causality for both traits

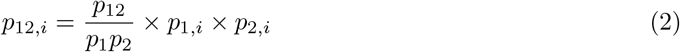

where *p*_12_, *p*_1_ and *p*_2_ are as described above. (See the Supplementary Text for more details.)

### Specification of variant-specific prior probabilities

Various methods are available for specifying variant-specific weights. In this paper we assessed four types of sources for specifying variant-specific weights (Table 1). First, we use prior probabilities of causality computed by the PolyFun method [13] applied to 19 million imputed UK Biobank SNPs with MAF > 0.1%, based on a meta-analysis of 15 UK Biobank traits. PolyFun is a method designed to compute prior probabilities in proportion to predicted per-SNP heritabilities given functional annotations. Second, we used the ‘Gnocchi’ score, a measure of non-coding genomic constraint estimated from 76,156 human genomes [11]. The idea of the score is that regions that show high constraint are more likely to have functional consequences; use of the Gnocchi score in fine-mapping of the UK Biobank association data increased posterior inclusion probabilities for a subset of variant-trait pairs [11]. Third, we used genome-wide enhancer-gene link scores calculated using the activity-by-contact (ABC) method [10, 19]. The ABC score measures the strength of enhancer-gene connections based on the activity of the enhancer and the contact between the gene and the enhancer. The contact is measured using genome-wide chromatin conformation capture techniques (Hi-C), that measure whether two DNA fragments physically associated in 3D space [20]. Finally, we used the empirical density of the distance of eQTLs to their associated TSSs we calculated from public datasets.

**Table 1:** Methods for specifying variant-specific prior probabilities

| Method | Used for fine-mapping? | Gene-specific? | Dataset | Citation |
| --- | --- | --- | --- | --- |
| PolyFun | Yes | No | NA | [13] |
| Gnocchi | Yes | No | NA | [11] |
| eQTL-TSS distance | No | Yes | eQTLGen<br>OneK1K round 1<br>OneK1K round 2 | - |
| Activity-by-contact (ABC) score | No | Yes | NA | [10, 19] |

To compute an empirical eQTL-TSS distance density we used several publicly available datasets: the datasets in the eQTL Catalogue [21], GTEx v8 (accessed through the eQTL catalogue) [22], eQTLGen [23] and the OneK1K dataset [24]. First, we filtered all the eQTL datasets to genome-wide significant eQTLs and the most significant eQTL for each gene, and then calculated the distance of each to eQTL to the TSS of the associated gene (Methods). To determine which datasets to estimate the density from we assessed the distribution of distances from the TSS across all the datasets (Figure S1). This analysis showed relatively little variation between tissues in GTEx or the eQTL catalogue, suggesting that tissue-specific densities are not needed. It also highlighted that eQTLs in the datasets eQTLGen and OneK1K were generally more distal from the TSS. Investigating the OneK1K dataset further, we found that eQTLs that were specific to a cell type (Figure S2c) and identified in subsequent rounds of conditional mapping (Figure S2a) were more distal [25] while distance was consistent across cell types (Figure S2b) except for a few rare cell-types. Therefore, based on evidence that GWAS variants lie further from gene TSSs than eQTL [8] and empirical evidence conditional estimation can lead to more colocalisations [25] we selected the eQTLGen, cell-type specific OneK1K round 1 of conditional and OneK1K (Round 2-5) as the being the most appropriate datasets to estimate prior information from. (Methods) For the PolyFun, Gnocchi and ABC score priors we accessed publicly available data and performed minor processing, mainly to account for SNPs which did not have specific prior values (Methods).

Overall, these findings gave us 6 sources of variant-specific prior probabilities, shown in Figure 1 for a ±500Kb window around the TSS of *VAV3*. The eQTL-TSS density priors place most mass in a small region around the TSS, with the OneK1K round 2+ estimated prior being less concentrated around the TSS. The ABC score-prior is also centred around the TSS, but is less smooth.

**Figure 1:**
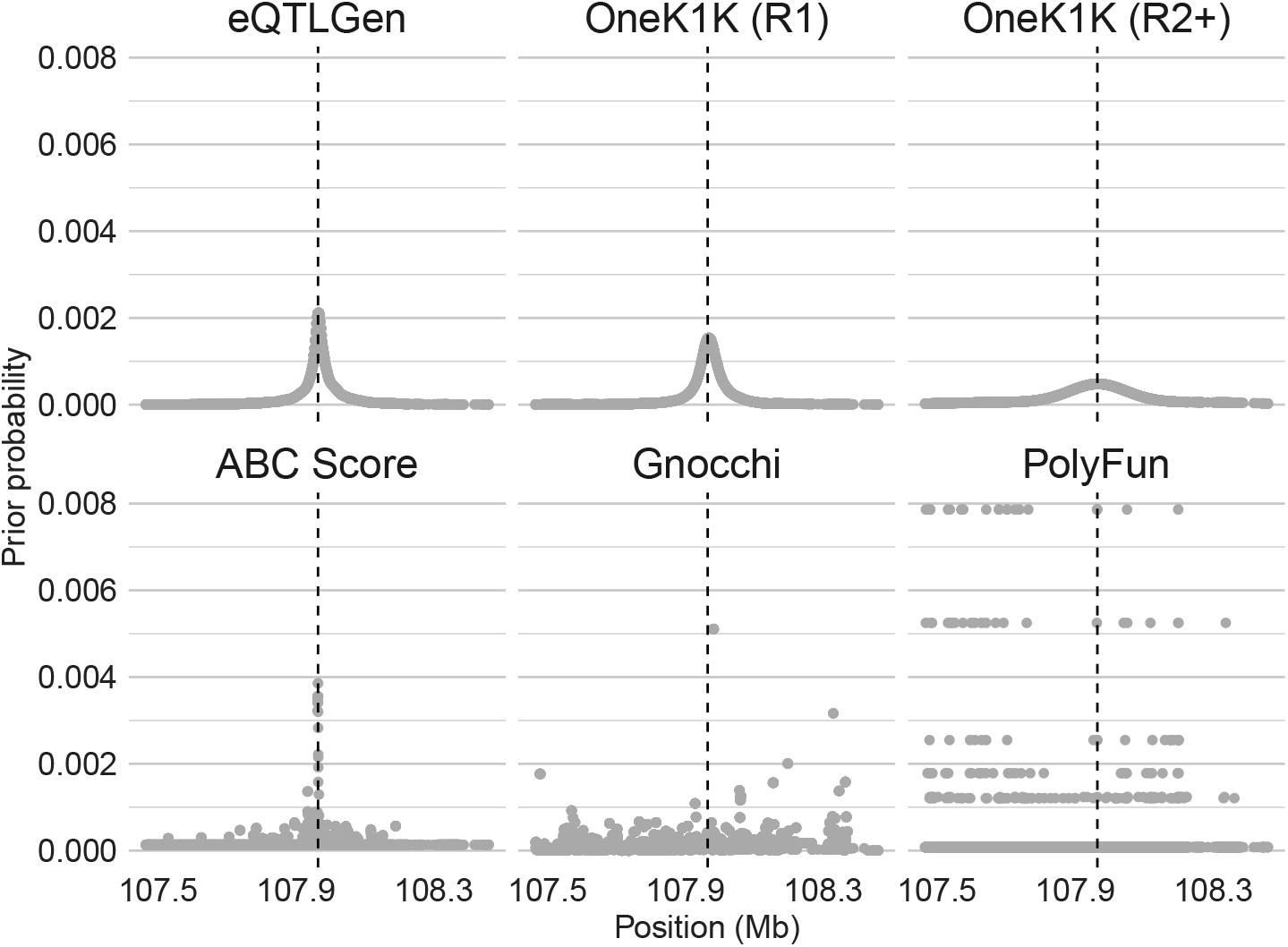
Prior probabilities of causality in a 1MB region around the TSS of *VAV3*. Prior probabilities from the six sources of prior probabilities: eQTLGen estimated eQTL-TSS distance density, OneK1K round 1 estimated eQTL-TSS distance density, OneK1K round 2+ estimated eQTL-TSS distance density, genome-wide enhancer-gene link scores calculated using the ABC score, Gnocchi non-coding constraint score and previously calculated PolyFun prior probabilities. The scores are displayed in a +/-500kB region around the canonical TSS, shown by the dashed black line, of the *VAV3* genes

### Simulation analysis

First, as a proof of concept, we performed simulations to validate our implementation and highlight the potential benefits of incorporating variant-specific prior information into colocalisation analysis. We simulated association summary statistics for ±500Kb regions around the TSS of three genes with patterns of high LD. High LD regions were chosen as high LD makes determining coloclaisation more challenging. Summary statistics were simulated using simGWAS [26] under both *H*_3_ and *H*_4_, using the eQTLGen-calculated distance density as the true density of causal variants (See Methods for details). Notably, the simulations recapitulate the tendency of Pr(*H*_4_) to be close to concentrated close to 0 or 1, for example as in the colocalisation calculated in the OpenTargets Genetics platform. (Figure S3b, see Methods for details). The original version of coloc, which assumes a single causal variant, was run on the simulated data, with and without using the simulation ground-truth distance density as the variant-specific prior.

Overall, using variant-specific priors improved the accuracy of colocalisation. Across all three loci, using variant-specific priors generally increased the computed Pr(*H*_4_) when the data was simulated under *H*_4_ and decreased it when the data was the simulated under *H*_3_ (Figure 2a). This finding demonstrates the ability of variant-specific priors to improve colocalisation accuracy, at least when the specified prior matches the true distribution of causal variants. In addition, we assessed how the effect of the variant-specific priors was effected by both the distance of the causal variant to the TSS and the value of Pr(*H*_4_) when uniform priors were used. We found that, as intended, variants, close the TSS had the largest increase in Pr(*H*_4_) while distal variants had smaller increases (Figure 2b). We also observed that simulations with moderate values of Pr(*H*_4_) using uniform priors had the largest change in Pr(*H*_4_) when variant-specific priors were used (Figure 2c). This observation is partly due to a ‘ceiling effect’: as 0 ≤ Pr(*H*_4_) ≤ 1, the many values close to 1 cannot be increased by a large amount by the variant-specific priors. The observation also highlights that variant-specific priors may only be able to change whether colocalisation is called at the widely used 0.8 threshold in the minority of cases where Pr(*H*_4_) is close to 0.8 using uniform priors.

**Figure 2:**
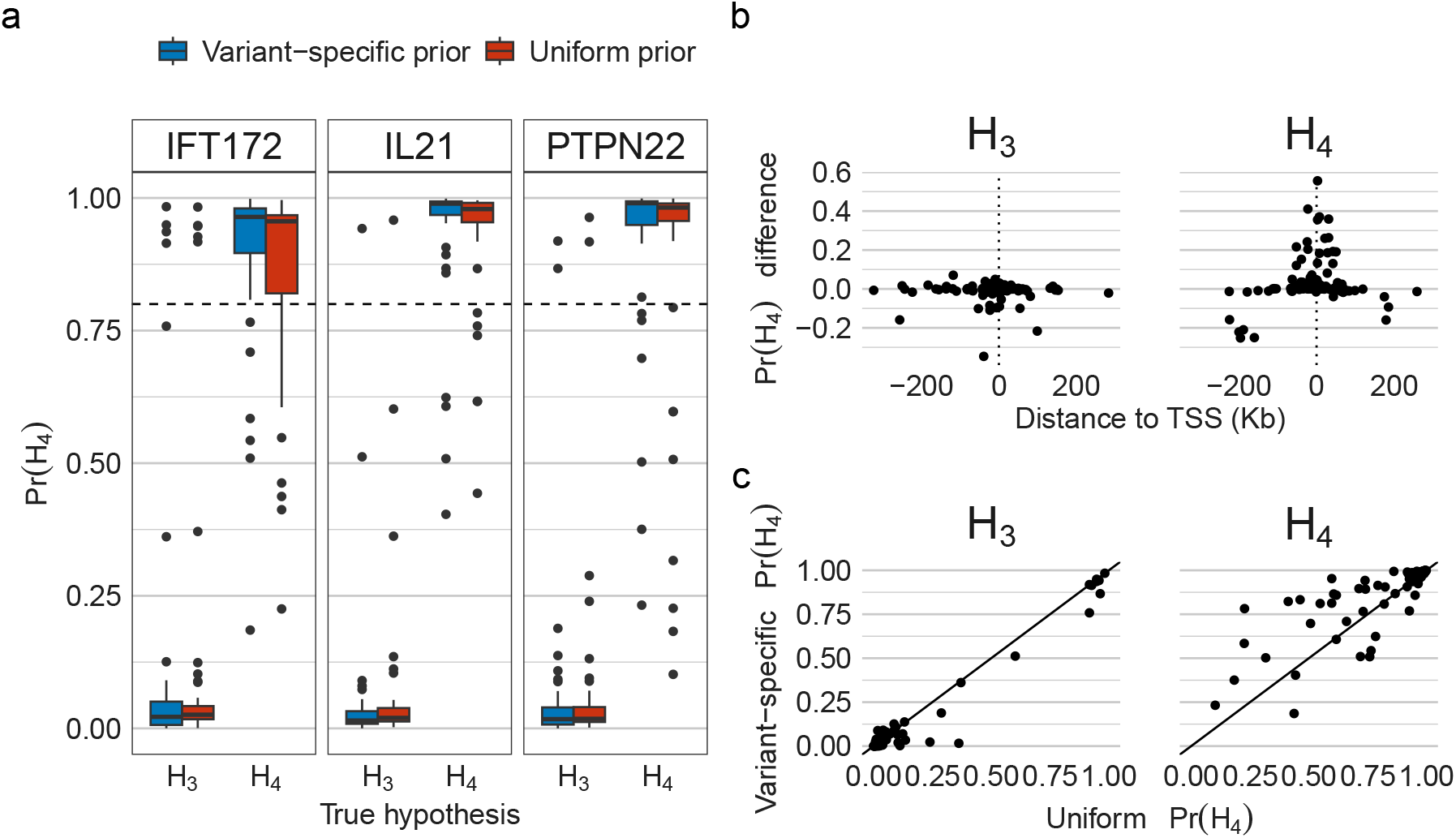
Simulation analysis. a) Pr(*H*_4_) values calculated using uniform priors or the simulation ground-truth distribution of causal variants as the variant-specific prior. Simulations are conducted under either coloc hypotheses *H*_3_ or *H*_4_ in 1MB (+/-500kb) regions around the TSS of the listed genes. b) For *H*_3_ and *H*_4_ simulations, the Pr(*H*_4_) calculated using variant-specific priors - Pr(*H*_4_) calculated using uniform priors against the position of the simulated causal variant. c) For *H*_3_ and *H*_4_ simulations, scatter plot of the Pr(*H*_4_) calculated using variant-specific priors against Pr(*H*_4_) calculated using uniform priors.

### pQTL-eQTL colocalisation performance comparison

Next, we sought to both evaluate the effectiveness of variant-specific priors in real world data and compare the performance of different sources of prior information. Comparing the performance of the prior-specification methods using simulations is challenging, because it is not clear how to realistically specify the true distribution of causal variant location without using information from prior sources. Instead, we compare the methods on their ability to recover ‘ground-truth’ pQTL-eQTL colocalisations, inspired by the comparison of colocalisation methods performed in [27]. The idea of this comparison is that we expect, based on the central dogma of molecular biology, that if a variant affects protein levels (i.e. is a pQTL) of a protein-coding gene then it should act by affecting that gene’s expression (i.e. is an eQTL). Therefore, the large majority of detectable cis-pQTLs should have a colocalising eQTL signal. We therefore compare the method’s ability to recover these ‘ground-truth’ pQTL-eQTL pairs for protein coding genes. Specifically, we colocalise pQTL for 3,215 proteins detected in blood plasma measured in 3,301 individuals from the INTERVAL study [28] against eQTL measured in five datasets from the eQTL catalogue (Table S1).

We applied all six prior-specification methods to each of the pQTL and eQTL data separately. In addition, for the eQTL TSS densities we applied it to both datasets simultaneously. Colocalisation was performed using both the original version of coloc, which assumes a single causal variant (coloc-single), and the version of coloc which uses SuSiE to relax that assumption (coloc-susie) (See Methods for details). The methods were run on 1Mb windows around the TSSs of protein-coding genes against 5 datasets from eQTL catalogue (Table S1, see Methods for more details). We applied the eQTL-TSS distance density priors centred at the TSS of the eQTL gene when applied to the eQTL dataset and centred at the TSS of gene associated with the protein applied to the pQTL dataset. We summarised the results of the analysis in two ways. First, we calculated the recall and precision of methods at the Pr(*H*_4_) > 0.8 threshold for colocalisation, calling colocalisation if the pQTL colocalised across any of the eQTL datasets (Figure 3a, Figure S4a). Second, we used a ROC-style analysis to assess the performance of including the prior information over a range of Pr(*H*_4_) thresholds (Figure 3b, Figure S4b). (See Methods for details).

**Figure 3:**
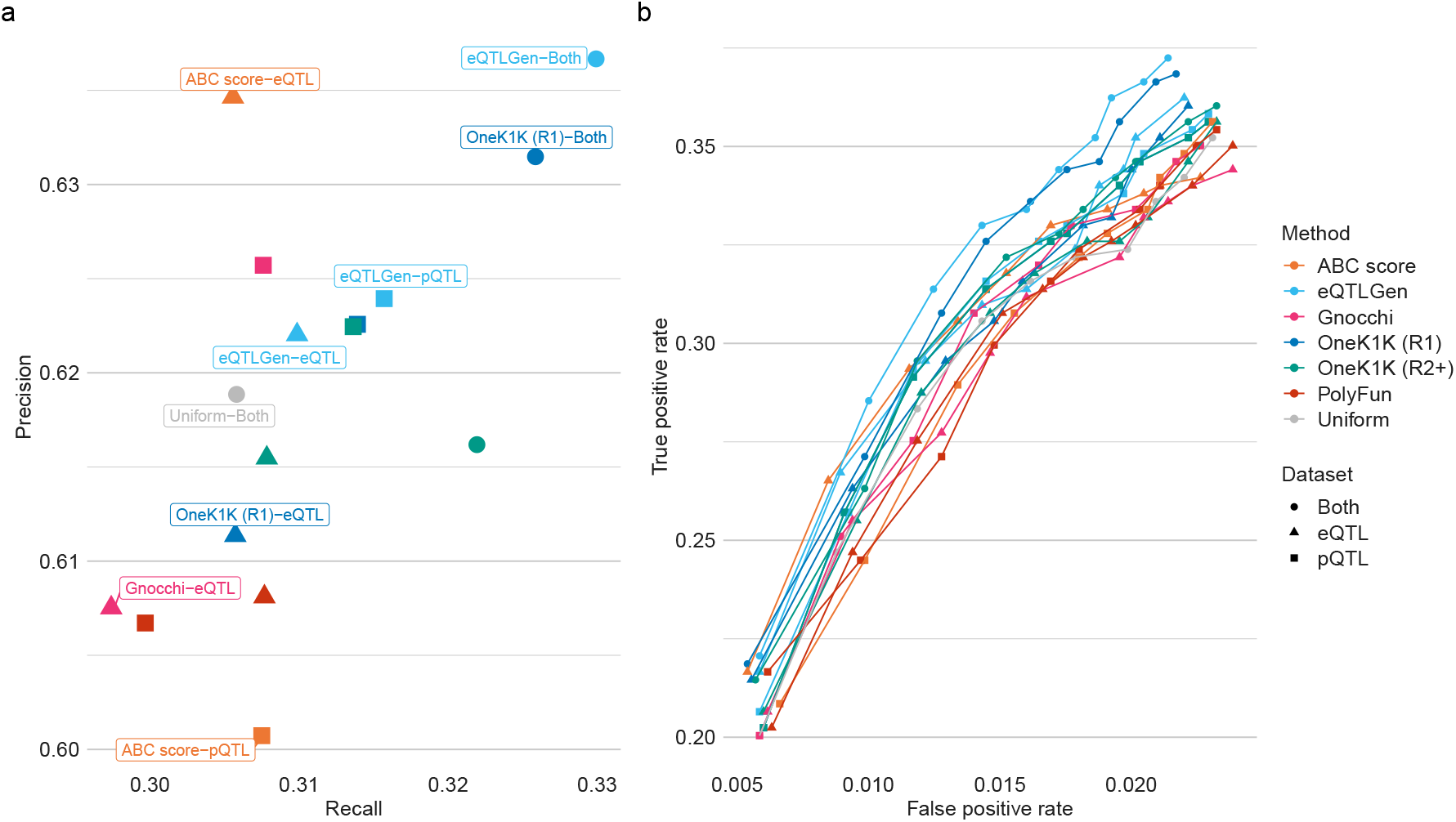
pQTL-eQTL colocalisation performance assessment for coloc with a single causal variant assumption. a) Recall and precision of colocalisation at the Pr(*H*_4_) > 0.8 significance threshold based on ground-truth pQTL-eQTL colocalisations. All 6 different sources of prior information were applied to either the eQTL or pQTL data, with the 3 eQTL-TSS distance density priors additionally applied to both datasets. b) A ROC-style analysis of the ground-truth pQTL-eQTL data using the Pr(*H*_4_) thresholds: 0.5, 0.55, …, 0.95. In both panels coloc assuming a single causal variant (coloc-single) is used.

There were three main findings from this analysis. First, the variant-specific priors had a positive but modest improvement in performance for both coloc-single and coloc-susie. At the 0.8 threshold, the maximum improvements in recall from incorporating prior information was 0.04 and the maximum improvement in precision was 0.03. Prior information had a larger effect on performance when Pr(*H*_4_) threshold was lower, corresponding to higher true and false positive rates. At the 0.8 threshold, all sources of prior information improved recall but at the cost of precision for the ABC score prior applied to the pQTL dataset, OneK1K Round 2 and Gnocchi priors applied to the eQTL dataset. Second, when applied to one dataset only, the best performing priors at the 0.8 threshold were the eQTLGen-estimated density prior applied to the eQTL and pQTL datasets, the OneK1K R1-estimated density prior applied to the pQTL dataset and the ABC score prior applied to the eQTL dataset. The worst performing priors were the PolyFun prior applied to the eQTL dataset, the OneK1K round 2-estimated prior applied to the eQTL dataset and the ABC score prior applied to the pQTL dataset. Across thresholds, the eQTL-estimated priors shows the best performance. Finally, we observed much better performance when the eQTL-estimated priors were applied to both pQTL and eQTL datasets simultaneously. This good performance makes sense as applying the priors to both datasets strongly encodes our knowledge that the pQTL and eQTL signals for the same gene/protein should be close. Furthermore, comparing results for coloc-single and coloc-susie, the different priors had similar relative effects on performance and the overall ordering of priors was the same for the two methods. However, coloc-susie had substantially higher recall but a slightly decreased precision (Figure S4).

To better understand the modest performance improvement of the priors in this comparison we assessed their overall impact on the posterior probability of colocalisation. We observed that all the priors had only a small effect on the calculated value of Pr(*H*_4_) (absolute value of change in Pr(*H*_4_) less than 0.01) in over 80% of loci. (Figure S5a). Using the variant-specific priors with coloc with SuSiE had a smaller effect on Pr(*H*_4_) across methods compared to coloc with the single causal variant assumption, particularly when applied to the eQTL datasets. We ascribe this difference to the higher maximum log Bayes factors calculated by SuSiE compared to calculated by the coloc-single method, suggesting that there is more information in the likelihood for coloc-susie (Figure S5b). Similarly, we observed that the magnitude of log Bayes factors were related to the size of the effect of variant-specific priors, with large effects only occurring when the maximum was small (Figure S5c). Focusing on the impact of the eQTLGen density prior, we observed that the values of Pr(*H*_4_) with uniform and variant specific priors, were highly correlated across loci, with only a minority of loci displaying large enough changes to alter conclusions at the 0.8 threshold (Figure S5b). Threshold effects explain some of the lack of changes, but they do not explain the absence of loci where the variant-specific prior substantially reduces Pr(*H*_4_) (Figure S5c). We hypothesised that the lack of changes, unlike in finemapping, could be due to colocalisation posterior probabilities being more concentrated towards 1 than probabilities produced by fine-mapping. However, comparing colocalisation posterior probabilities and SuSiE fine-mapping causal probabilities in the OpenTargets catalogue, indicated that both types of probabilities are highly concentrated close to 1 (Figure S3).

Overall, this analysis showed that incorporating prior information improves colocalisation performance in real colocalisations but the magnitude of the improvement is modest. The best performing source of prior information were the empirical eQTL-TSS distance densities.

### GWAS-eQTL colocalisations analysis

Finally, we assessed the impact of variant-specific priors on GWAS-eQTL colocalisation in order to understand how colocalisation results might change with the use of distance priors on eQTL studies. We performed colocalisation between three FinnGen (Release 10) GWAS (Autoimmune disease, Type 2 diabetes, and hypertension) and five eQTL datasets for disease-relevant tissues for each trait drawn from the eQTL catalogue (Table S2, see Methods for details). The colocalisation results using a uniform prior were compared to those using the eQTLGen and OneK1K round 1-estimated eQTL–TSS distance priors, which had the best performance in the pQTL–eQTL comparison.

Across trait-dataset pairs the variant-specific priors had a moderate impact on changing whether a variant-gene pair was significant (Figure 4a), with at most 13.5% of colocalisations with some evidence of colocalisations (Pr(*H*_4_) > 0.5)) changing significance at the 0.8 threshold. Overall, less colocalisations changed when the priors were used with coloc with SuSiE compared to coloc with the single variant assumption, while the different eQTL-TSS distance priors had very consistent effects. Examining the effect of priors on Pr(*H*_4_) showed that the for most loci the priors had little effect (Figure 4b), particularly for coloc with SuSiE (Figure 4c). Colocalisations that changed significance for either coloc-single (Supplementary Table 1) or coloc-susie (Supplementary Table 2) are recorded in the supplementary materials.

**Figure 4:**
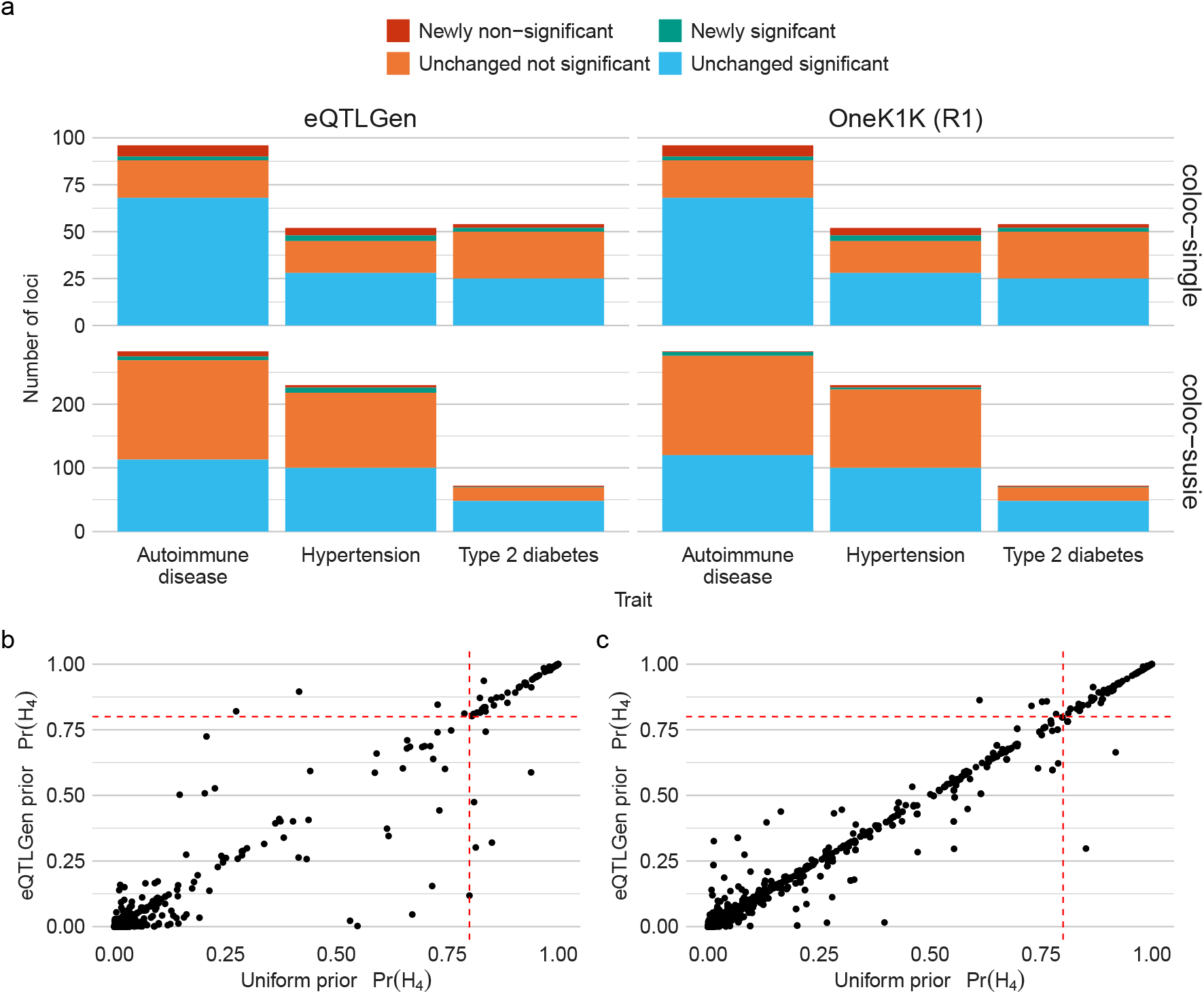
GWAS-eQTL colocalisation analysis. a) Number of loci with some evidence of colocalisation (Pr(*H*_4_) > 0.5) using a uniform prior coloured by the effect of a variant-specific priors on colocalisation significance threshold of 0.8. Results are for the eQTLGen and OneK1K round 1-estimated priors and coloc using SuSiE and coloc with the single variant assumption. Unlike in the pQTL-eQTL analysis, here colocalisations with each credible set are counted as separate colocalisations. b) Scatter plot of Pr(*H*_4_) computed using uniform and eQTLGen eQTL-TSS distance density priors for coloc with the single causal variant assumption. c) As in b) but using coloc with SuSiE.

One GWAS signal for which using variant-specific priors changed colocalisation conclusions was a FinnGen automimmune disease trait signal near the genes *PSMB7* and *NEK6*. Using coloc-single, this signal strongly colocalised with eQTL associations for both *PSMB7* and *NEK6* in GTEx thyroid tissue (Figure 5d). Extended LD means the GWAS signal is spread across both genes although the peak along this “plateau” is nearer the NEK6 TSS than the PSMB7 TSS. In the eQTL data, the peak of each gene signal is close to their respective TSSs. (Figure 5a, b, c), When the eQTLGen prior, centred at the *NEK6* TSS, is applied to the GWAS-*NEK6* colocalisation it remains significant (Pr(*H*4) = 0.98), but when the prior, centred at the *PSMB7* TSS, is applied to the GWAS-*PSMB7* colocalisation its Pr(*H*_4_) decreases to 0.42. Therefore, incorporating prior information at this locus changes the conclusion of colocalisation to strongly prefer *NEK6* as the causal gene. This example highlights the ability of variant-specific priors to resolve multiple colocalisations at a locus in a natural, distance-dependent way.

**Figure 5:**
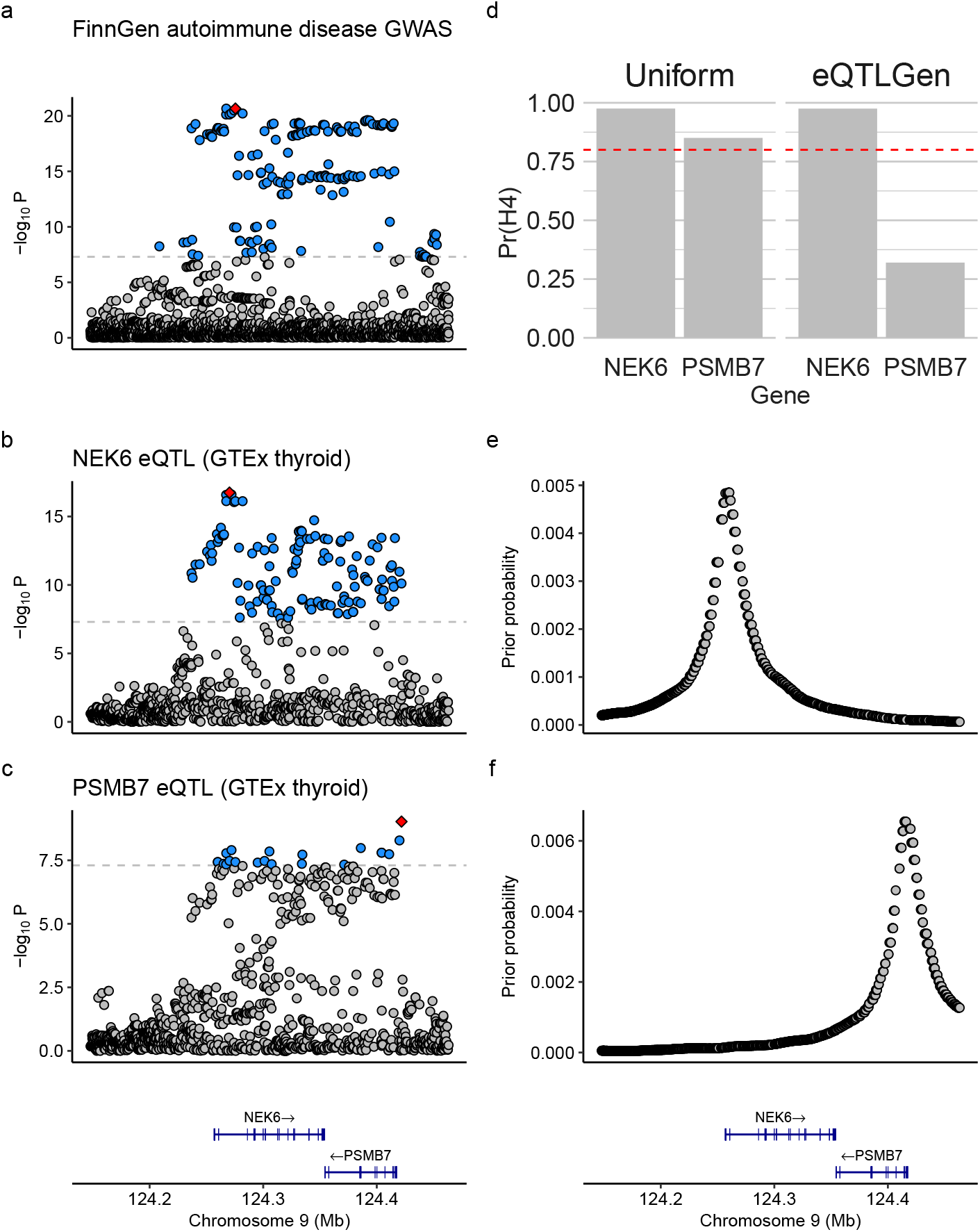
Colocalisation at the *NEK6*-*PSMB7* locus. a) Manhattan plot of the FinnGen v10 ‘Autoimmune disease’ trait GWAS in a ±500Kb region around the TSS of *NEK6* b) Manhattan plot of the *NEK6* GTEx thyroid eQTL results in the same region. c) Manhattan plot of the *PSMB7* GTEx thyroid eQTL results in the same region including gene track information showing *NEK6* and *PSMB7* d) Pr(*H*_4_) returned by coloc for colocalisation between the ‘Autoimmune disease GWAS’ and the *NEK6* and *PSMB7* eQTL results using both uniform priors and eQTLGen-estimated eQTL-TSS distance priors for each genes. e) The eQTLGen-estimated eQTL-TSS distance prior probabilities for *NEK6* f) The eQTLGen-estimated eQTL-TSS distance prior probabilities for *NEK6* with the same gene track as in c).

## Discussion

In this paper we described an implementation of variant-specific prior probabilities in the widely-used coloc method for statistical colocalisation. Our simulations and pQTL-eQTL colocalisation analysis demonstrated that incorporating prior information can improve the accuracy of colocalisation. However, the improvements were modest, with 2-3% improvements in recall and precision in comparisons based on ground truth pQTL–eQTL colocalisations. In this context, pQTL–eQTL-based comparisons are very useful to compare the effect of varying priors for coloc, and there is no equivalent ground truth dataset in fine-mapping. However, we note that our analysis is likely to overestimate false negatives, because the proteins in plasma originate from a wide array of tissues and our eQTL datasets cover only a subset of these. Overall, the best performing source of prior information were the estimated eQTL-TSS distance densities. In GWAS-eQTL colocalisations, using the eQTL-TSS density as source of variant specific prior information changed colocalisation significance in up to 13.5% of all loci with some evidence of colocalisation (Pr(*H*_4_) > 0.5). The eQTL-TSS density priors could sensibly distinguish between multiple gene with significant colocalisations, for example at the *NEK6* /*PSMB7* locus.

In contrast to previous reports in fine-mapping [13], all variant-specific priors did not alter conclusions in a large number of cases. We speculate this lack of effect is due to fine-mapping being more sensitive to priors probabilities than colocalisation. For example, a fine mapping prior might only need to up-weight one variant to change a credible set of size two to size one and clarify the causal variant. However, to affect the probability of colocalisation, which is computed by summing across variants, we likely need to change the posterior probabilities for the majority of variants that provide evidence of colocalisation. This interpretation is supported by the greater impact of the eQTL-TSS distance density priors, which consistently alter prior probabilities for a large proportion of variants at a locus, compared to other sources of prior information. However, we note our findings are consistent with the reported effect of using the Gnocchi score to specify variant-specific priors for fine-mapping where only 0.3% of variants were newly identified as likely causal (PIP > 0.8) [11]. In addition, in our comparisons the variant-specific priors, other than the eQTL-TSS density distance priors, did not consistently improve eQTL-pQTL colocalisation analysis. This lack of improvement may be because the priors do not capture of position of causal eQTL variants, for example because of systematic differences between eQTLs and GWAS hits [8]. Our comparison highlights the lack of systematic comparison of different strategies for setting prior probabilities in statistical fine-mapping.

In this paper we incorporated additional information at the variant level in colocalisation analysis by altering prior probabilities. The limited impact of this approach suggests that other methods of incorporating additional information may be preferable. One approach would be to introduce additional information at the GWAS analysis stage, capturing that information in updated Bayes factors that could be used by coloc. Another approach is to incorporate additional information after colocalisation has been performed, as in locus-to-gene models such as OpenTarget’s L2G model [15], which use colocalisation probability as a covariate along with information such as distance and predicted variant effect.

The ability to use variant-specific priors is currently available in the variant-specific-priors branch of coloc and will be included in the forthcoming v6.0.0 release of coloc. Based on this analysis conducted in this paper, we recommend that eQTL-TSS distance densities priors be strongly considered for use in all GWAS-eQTL colocalisations. In our comparisons, other sources of prior information did not have a large enough impact to justify their use. However, our results do not preclude the use of additional bespoke sources of information, such as readouts from disease relevant cell-types, which we did not explore due to a lack of ground-truth benchmark. Any future sources of prior information that are demonstrated to improve fine-mapping results could be tried in colocalisation analysis. We expect that variant-specific priors will also be useful to better understand colocalisations of interest, such as colocalisations that show borderline significance. Practically, we advise that variant-specific priors should only be applied to one trait at a time, unless there is strong independent prior information for both traits. (See the Supplementary Text for more discussion of applying priors to both traits.) Finally, we emphasise that any bespoke variant-specific priors used in an analysis should be transparently reported and explained, and their use justified in detail.

Finally, this paper has a two main limitations. First, for computational reasons we did not directly apply PolyFun to the GWAS traits of interest. The PolyFun method advises that PolyFun should be trained on each GWAS dataset of interest, but this is computationally expensive, especially when we compared multiple pQTL and eQTL datasets. The PolyFun software provides pre-computed priors averaged over multiple UK Biobank traits, which we used, but an individually trained prior may have performed better. Second, the pQTL-eQTL results of colocalisation comparison may not perfectly generalise to the GWAS-eQTL setting, due to the differences in genetic architecture between e/pQTL and GWAS hits [8].

## Methods

### eQTL data processing

To estimate the eQTL-TSS distance density from summary statistics, we used the following strategy. First, we filtered the eQTL summary statistics to only include genome-wide significant (*p* < 5 × 10^−8^) eQTLs, and only the most significant eQTL for each gene. (This approach was inspired by [29]) Information about the TSS and strand of each gene was downloaded from Ensembl with the biomaRt R package [30]. If a gene had multiple listed TSSs the median of their locations was used as the TSS. For each eQTL with position *p* say, the distance, *d* to the TSS, with position *t* say, was calculated as

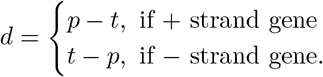

### Prior probability sources

#### ABC score

The ABC score for a gene *G* contributed by element *E* is,

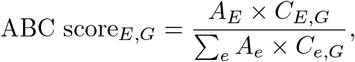

where *A*_*E*_ is the activity, its strength as an enhancer, of element *E* and *C*_*E,G*_ is the Hi-C measured contact between element *E* and gene *G*. The sum in the denominator is taken over all elements within 5 Mb of *G* [19].

We use ABC data collected for 131 biosamples generated by [10], which includes details of how activity and contact are measured. The dataset only includes gene-element connections with an ABC score ≥ 0.015. To convert the dataset into a vector of prior probabilities we filter to the gene of interest, take the median over all biosamples and set the value for a SNP to the value of the enhancer it lies in. If the SNP does not have a listed score we set the value to 0.0075, half the minimum score.

#### PolyFun

PolyFun calculates prior causal probabilities that are proportional to the ‘per-SNP heritability’ estimates, var(*β*_*i*_ | ***a***_*i*_), that is it sets,

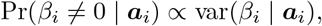

where *β*_*i*_ is the effect size for SNP *i* and ***a***_***i***_ is the vector of functional annotations for SNP *i*. This relationship can be derived under a point-normal prior model for *β*_*i*_ and assuming that the causal variance is independent of ***a***_***i***_; that is that var(*β*_*i*_ | *β*_*i*_ ≠ 0, ***a***_*i*_) = var(*β*_*i*_ | *β*_*i*_ ≠ 0) [31].

To use the PolyFun weights we downloaded the weights pre-computed using 9 million imputed UK Biobank SNPs with MAF> 0.1%, based on a meta-analysis of 15 UK Biobank traits. We reimplemented the processing performed in `polyfun/extract snpvar.py` to link the weights to data. SNPs that did not have a variance entry were assigned the minimum entry present in the data.

#### Gnocchi

Gnocchi is a signed score that quantifies the depletion of genomic variation at 1Kb scale [11]. Precisely, the score is defined as,

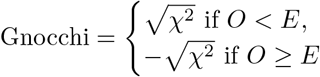

where *χ*^2^ = (*O − E*)^2^*/E* and *E* and *O* are the observed and expected values of the variation, respectively. The expected amount of variation is calculated based on estimated probabilities of mutations in trinucleotide contexts and adjusted for DNA methylation and genomic features. These calculations used data on 76,156 human genomes from gnomAD.

To assign variants to Gnocchi score we used the following approach. If a variant lay in a region with a Gnocchi score it was assigned that score. Otherwise it was assigned the score of the closest region with a Gnocchi score. The assigned scores were then converted to non-negative weights by a softmax transformation.

#### eQTL-TSS distance density

To estimate the densities we fit a density to the filtered eQTL’s distances using the R density() function. Prior probabilities for variants at a specific locus are then calculated using the density; SNPs are assigned the density value of the computed density point closest to the SNPs distance to the TSS. In the OneK1K dataset we use eQTLs that are detected in only 1 cell type. We filter eQTLs as either being from round of conditional testing 1 or round 2-5 and estimate two densities. For the eQTLGen dataset we estimate the density from all eQTLs.

To compute the density from the calculated distances we used a non-parametric approach to ensure maximum fidelity. We used the `density()` function in R, with non-default parameters `bw = “SJ”, cut = 0, adjust = 8`. The parameter `bw = “SJ”` specifies an alternate bandwidth selection algorithm that is recommended in the R documentation. The parameter `cut` specifies how far beyond the extremes of the data the density should become approximately 0. In practice, we found that values of cut > 0 produced a small number of points with much smaller values compared to the rest of the estimated density. The parameter `adjust` specifies a multiplicative scaling factor for the bandwidth. We set `adjust = 8` to heuristically account for uncertainty in the location of the TSS. The estimated density is represented computationally as 512 (distance, density value) pairs. The densities were computed in a *±*500kb window around the TSS.

### Simulation analysis

We simulated GWAS summary data under a single-causal variant assumption, simulating data under both *H*_3_ and *H*_4_. We used haplotypes for EUR samples in the 1000 Genomes phase 3 data [32], phased by IMPUTE2 [33], (https://mathgen.stats.ox.ac.uk/impute/1000GP_Phase3.html)

We used the `simGWAS` R package method to simulate GWAS summary statistics with the LD and MAF calculated from the haplotypes [26]. We simulated case-control data with 2, 000 cases and 2, 000 controls. The effect size of the causal variant was simulated as the maximum of 100 *N* (0, 0.0025) random variables. In each call to `simGWAS()` the simulation was repeated 2000 times and only the first simulation that had a minimum p-values less than 5 ×10^−6^ was used, matching our expectation that colocalisation is only performed when there is at least a moderate signal of association. Specifically we simulated data for ±500Kb windows around the TSS of three genes: *IL21, PTPN22, IFT172*. To simulate variant-specific priors we used the following algorithm. First, we sample the causal variant for the first trait from the eQTLGen density variant-specific prior. Second, sample the causal variant for the second trait, depending on the hypothesis:

- *H*_3_: sample the causal variant uniformly from all variants other than the causal variant for trait 1 or,
- *H*_4_: set the casual variant to the causal variant for trait 1.

### pQTL-eQTL colocalisation analysis

We used the pQTL dataset produced by mapping QTLs for 3,215 measured proteins in blood plasma from 3,301 individuals who were part of the INTERVAL study [28]. We colocalised the dataset against 5 eQTL datasets from the eQTL Catalogue (Table S1) chosen to maximise the number of colocalisations with plasma pQTLs. Summary statistics and fine-mapping credible set and log Bayes factor (LBF) files were downloaded from the eQTL Catalogue.

Using `coloc.abf()`, we performed colocalisation in a 1MB window around all protein-coding genes. Colocalisation was only performed if the region contained at least one pQTL with p-value < 5 × 10^−8^ and at least one eQTL with p-value < 5 × 10^−6^. The `coloc.abf()` function was run prior probabilities: *p*_1_ = *p*_2_ = 10^−4^, *p*_12_ = 5^−6^.

Using `coloc.susie()`, we performed colocalisation in a 1MB window around all protein-coding genes. Colocalisation was only performed for credible sets in both pQTL and eQTL datasets. The `coloc.bf_bf()` function was run with default prior probabilities (*p*_1_ = *p*_2_ = 10^−4^, *p*_12_ = 5 × 10^−6^)

For both approaches a colocalisation was defined as significant if the maximum colocalisation across eQTL datasets was Pr(*H*_4_) > 0.8. In the coloc with SuSiE analysis the maximum colocalisation was also taken across credible set pairs. To assess the performance of the methods we calculated various measures of classifier performance. To calculate these metrics we defined true positive, false positives, true negatives and false negatives in the following way,

- True positive: significant colocalisation between pQTL and eQTL for the same gene.
- False positive: significant colocalisation between pQTL and eQTL for different genes.
- False negative: no significant colocalisations but pQTL and eQTL genes match.
- True negative: no significant colocalisation and pQTL and eQTL genes do not match.

These metrics are only calculated for regions where colocalisation is performed i.e. where at least one pQTL with p-value < 5 × 10^−8^ and at least one eQTL with p-value < 5 × 10^−6^.

First, following the analysis in [27] we calculated the recall and precision of each prior method. These values were

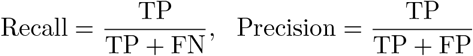

where TP is the number of true positives, FN is the number of false negatives and FP is the number of false positives. Second, we compute a receiver operator curve (ROC) for each method, for a series of colocalisation significance thresholds (0.5, 0.55, …, 0.95) for calculating the true positive rate (TPR) and false positive rate (FPR). These metrics are calculated as

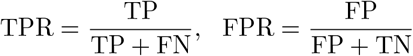

where FP is the number of false positives, FN is the number of false negatives and FN is the number of false negatives.

### GWAS-eQTL colocalisation analysis

Colocalisation was performed for three FinnGen traits (`AUTOIMMUNE, T2D WIDE, I9 HYPTENS`) against eQTLs in disease-relevant tissues (Table S2). We used GWAS summary statistics and fine-mapping credible set and log Bayes factor (LBF) files from FinnGen Release 10. In this analysis colocalisations with a signal captured by a single credible set are counted separately, unlike in the pQTL–eQTL analysis where the maximum is taken over colocalisations with all credible sets in a locus.

Using `coloc.abf()`, we performed colocalisation in a 1MB window around all protein-coding genes. Colocalisation was only performed if the region contained at least one genome-wide significant pQTL and at least eQTL with p-value less than 5 × 10^−6^. The `coloc.abf()` function was run with prior probabilities: *p*_1_ = *p*_2_ = 10^−4^, *p*_12_ = 5 × 10−6.

Using `coloc.bf_bf()`, was performed for overlapping GWAS and eQTL defined regions where there were credible sets for both GWAS and eQTL datasets. ‘Low purity’ credible sets present in the FinnGen data were removed. The `coloc.bf_bf()` function was run with default prior probabilities: *p*_1_ = *p*_2_ = 10^−4^, *p*_12_ = 5 × 10^−6^.

### Open Targets Genetics data

We downloaded the latest tranche of the Open Targets Genetics data from the Open Targets Genetics FTP site (https://ftp.ebi.ac.uk/pub/databases/opentargets/genetics/). We extracted the Pr(*H*_4_) from all the colocalisation results present in the v2d coloc folder and then filtered to Pr(*H*_4_) > 0.5. As analysing all variants was computationally impractical we randomly sampled 1,000,000 SuSiE results for variants from the v2d credset folder. The variants were filtered to be the lead variant in each locus and the posterior inclusion probability (PIP) was extracted. The PIPs were further filtered to be > 0.5.

## Data availability

The eQTLGen data was downloaded from https://molgenis26.gcc.rug.nl/downloads/eqtlgen/cis-eqtl/2019-12-11-cis-eQTLsFDR0.05-ProbeLevel-CohortInfoRemoved-BonferroniAdded.txt.gz. The OneK1K data was downloaded from https://onek1k.s3.ap-southeast-2.amazonaws.com/esnp/esnp_table.tsv.gz. The eQTL catalogue v6 data (including GTEx v8) was downloaded from the eQTL Catalogue FTP site using the links specified in https://raw.githubusercontent.com/eQTL-Catalogue/eQTL-Catalogue-resources/master/tabix/tabix_ftp_paths.tsv. The FinnGen GWAS summary statistics were downloaded following the FinnGen consortium’s instructions https://www.finngen.fi/en/access_results. The Open Targets Genetics data was downloaded from the Open Targets Genetics FTP site https://ftp.ebi.ac.uk/pub/databases/opentargets/genetics/. The 1000 Genomes phase 3 haplotype data was downloaded from https://mathgen.stats.ox.ac.uk/impute/1000GP_Phase3.html.

## Code availability

All code to reproduce the analyses presented in this paper, including a Snakemake pipeline [34], is available in a GitHub repository: https://github.com/jeffreypullin/coloc-variant-specific-priors/tree/main. The coloc package with support for variant-specific priors is available in the variant-specific-priors branch in the coloc repository https://github.com/chr1swallace/coloc/tree/variant-specific-priors and will be made generally available as part of coloc v6.0.0.

## Acknowledgements

We thank members of the Wallace group and the MRC Biostatistics Unit for helpful discussions. We acknowledge the participants and investigators of the FinnGen study.

## Supplementary figures and tables

**Supplementary Table S1:**
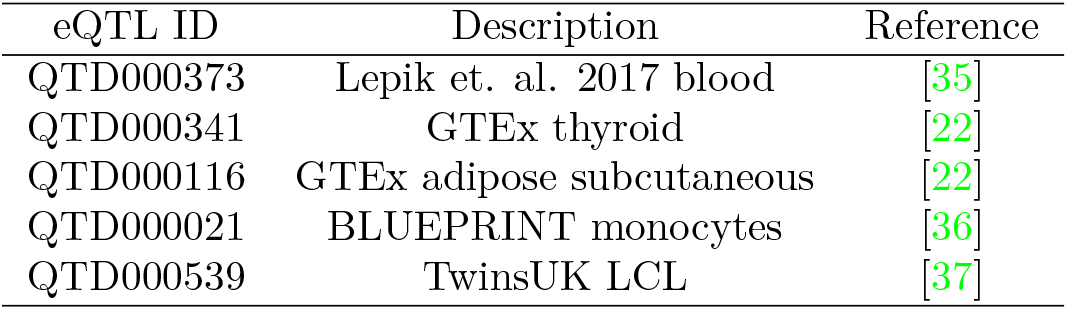
eQTL datasets used for pQTL-eQTL colocalisation

**Supplementary Table S2:**
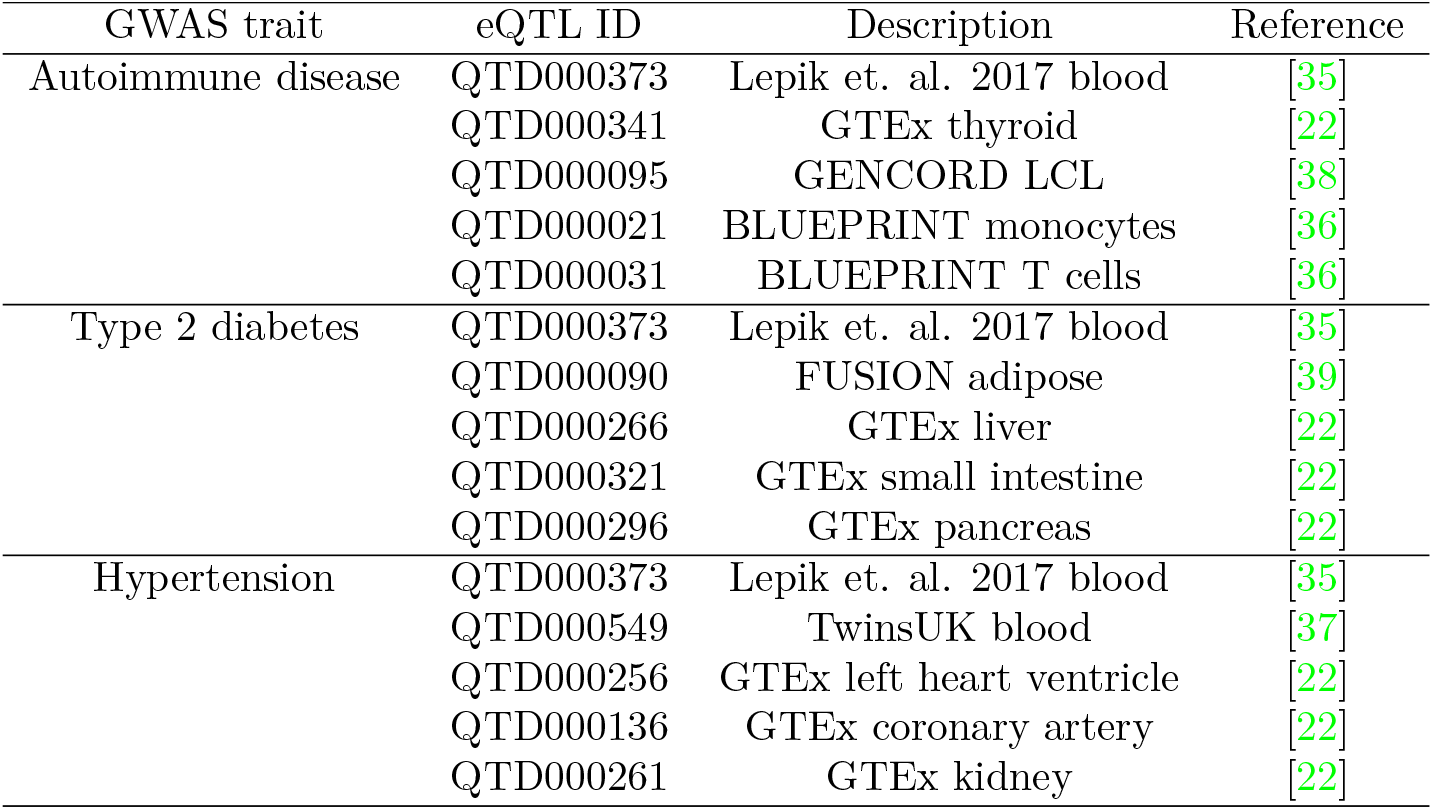
eQTL datasets used for GWAS-eQTL colocalisation GWAS trait eQTL ID Description Reference

**Supplementary Figure S1:**
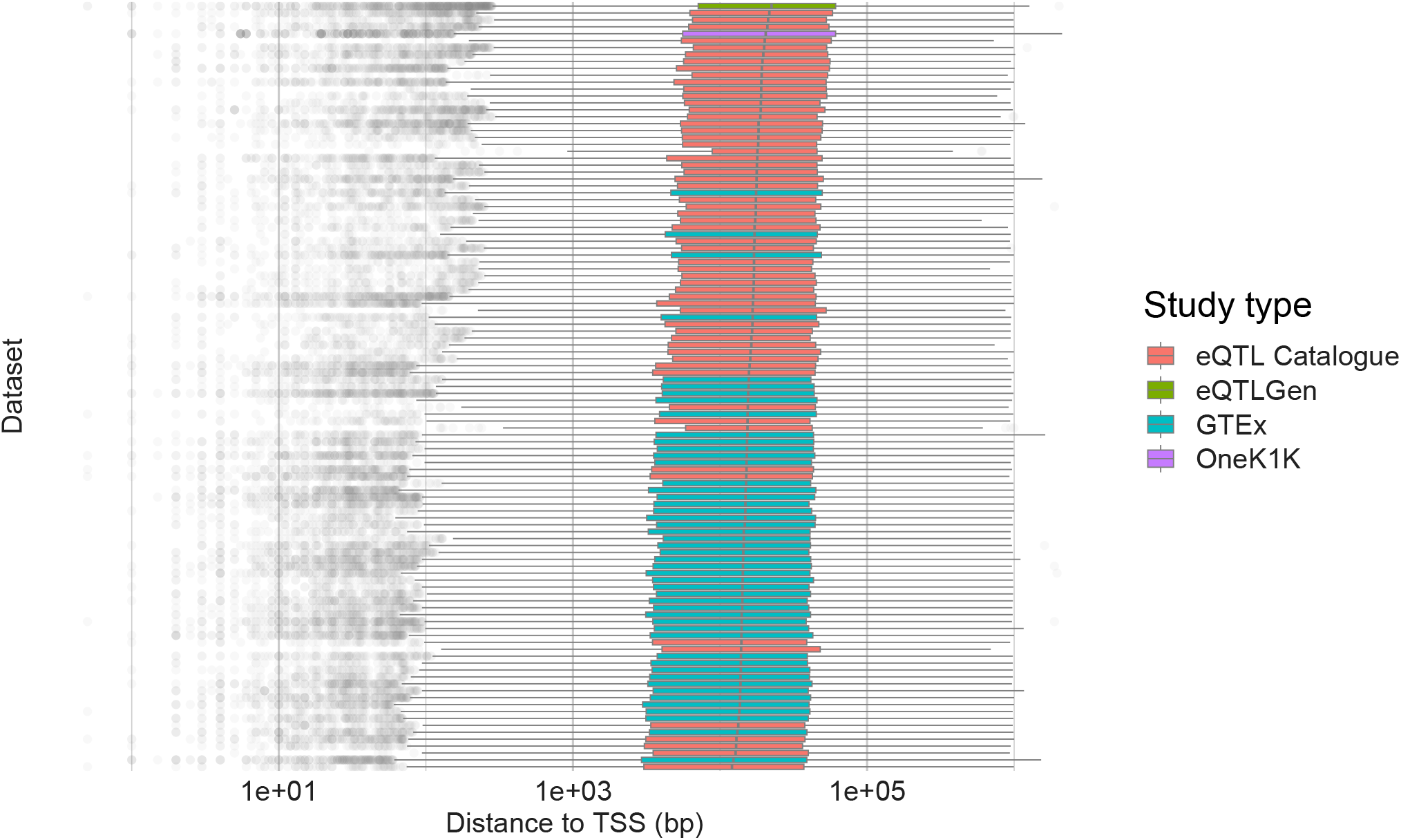
Distribution of eQTL-TSS distance across datasets. Boxplots of distance (measured in base pairs, log10 scale) of eQTLs to TSS for genome-wide significant eQTLs in v6 of the eQTL catalogue (including GTEx v8), eQTLGen and OneK1K dataset. Each boxplot corresponds to a different study.

**Supplementary Figure S2:**
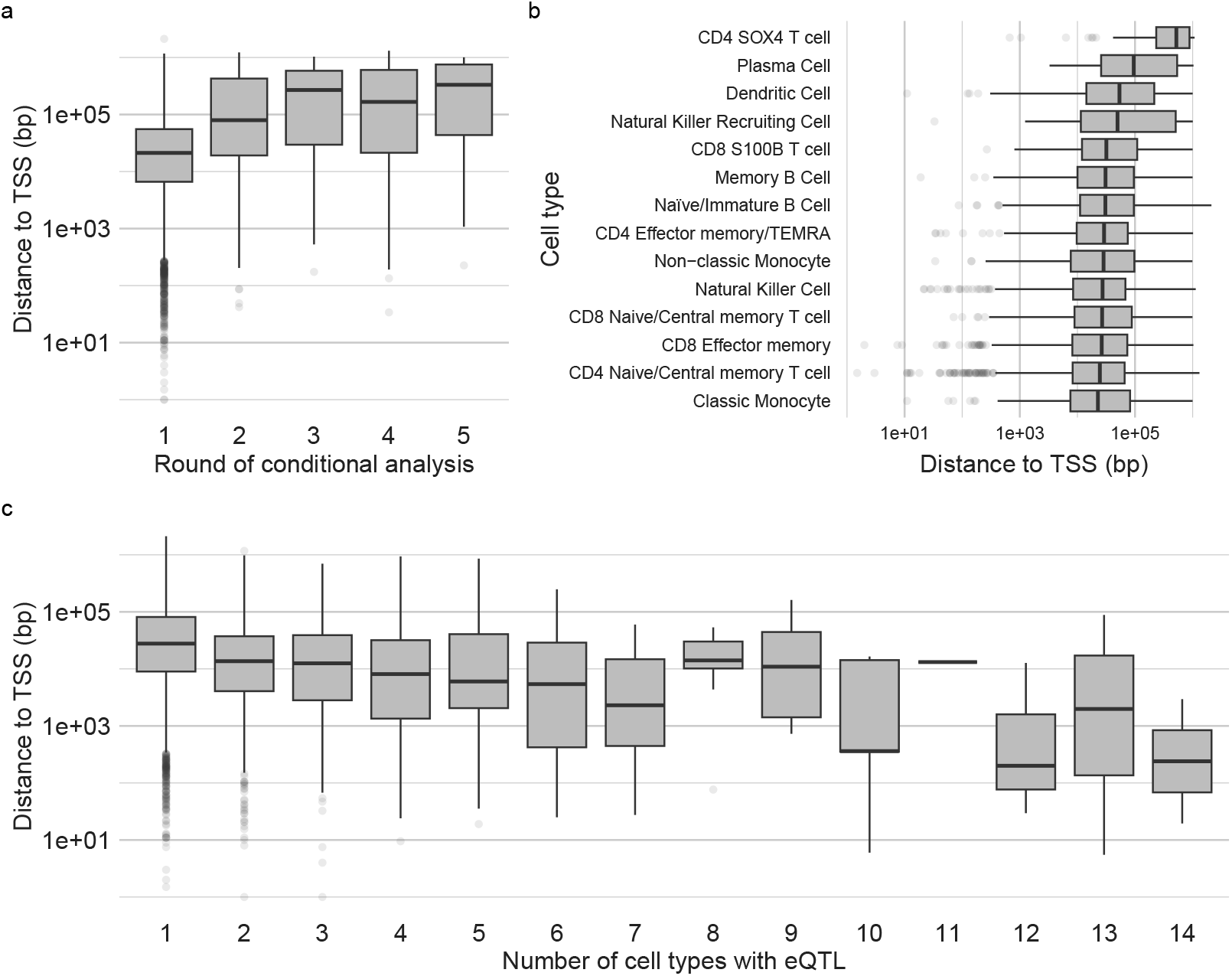
Distribution of eQTL-TSS distance in the OneK1K dataset. a) Boxplots of distance (measured in base pairs) of eQTLs to TSS (log10 scale) by round of conditional analysis. b) Boxplots of distance (measured in base pairs) of eQTLs to TSS (log10 scale) by cell type. c) Boxplots of distance (measured in base pairs) of eQTLs to TSS (log10 scale) by the number of cell types the eQTL is significant in.

**Supplementary Figure S3:**
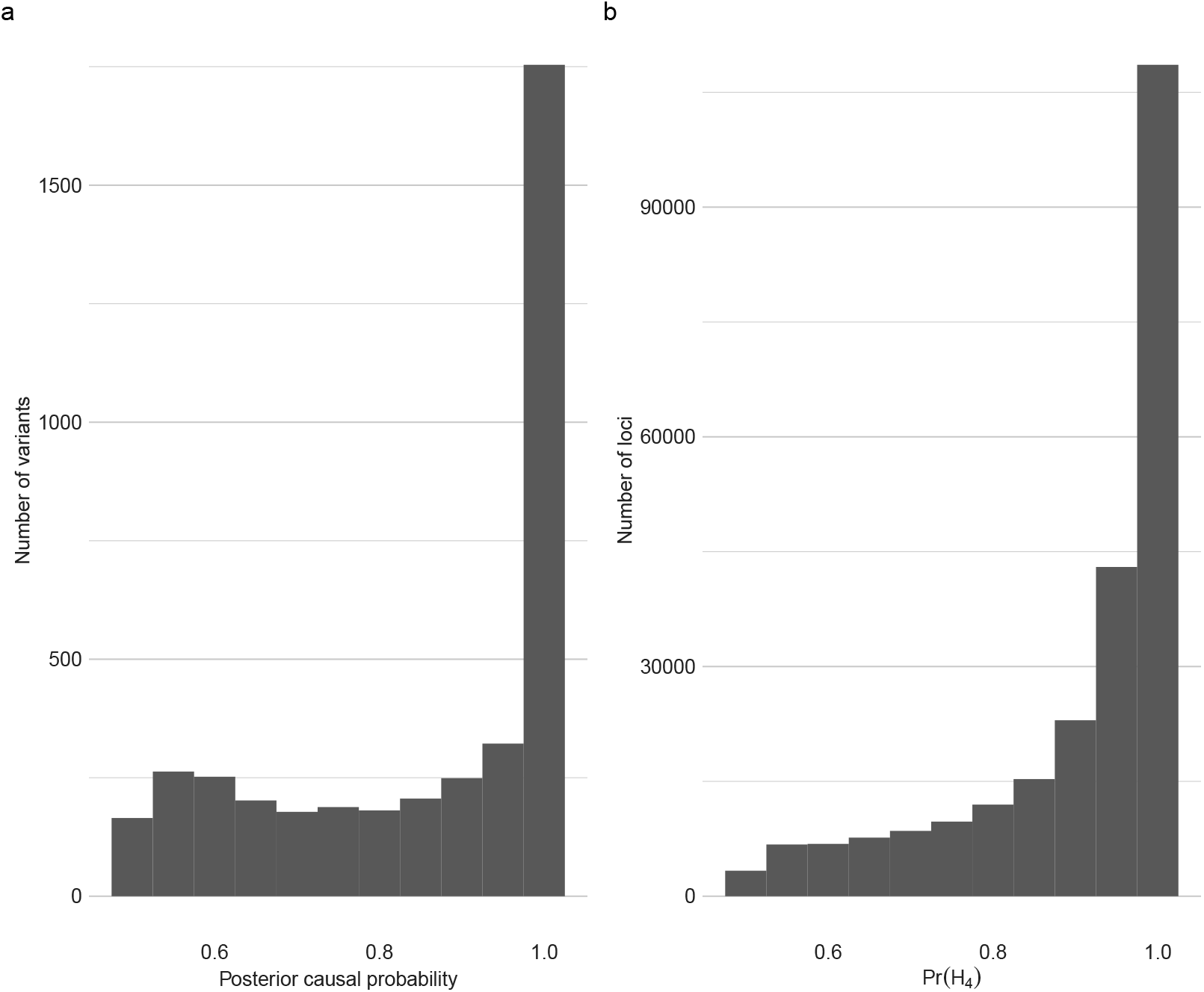
Credible set and colocalisations posterior probabilities in Open-Targets Genetics data. a) Random sample of posterior inclusion probabilities from the lead variant in 95% credible sets calculated using SuSiE in the OpenTargets Genetics platform b) All Pr(*H*_4_) calculated by coloc in the OpenTargets Genetics platform. Both plots are restricted to probabilities > 0.5. (See Methods for details.)

**Supplementary Figure S4:**
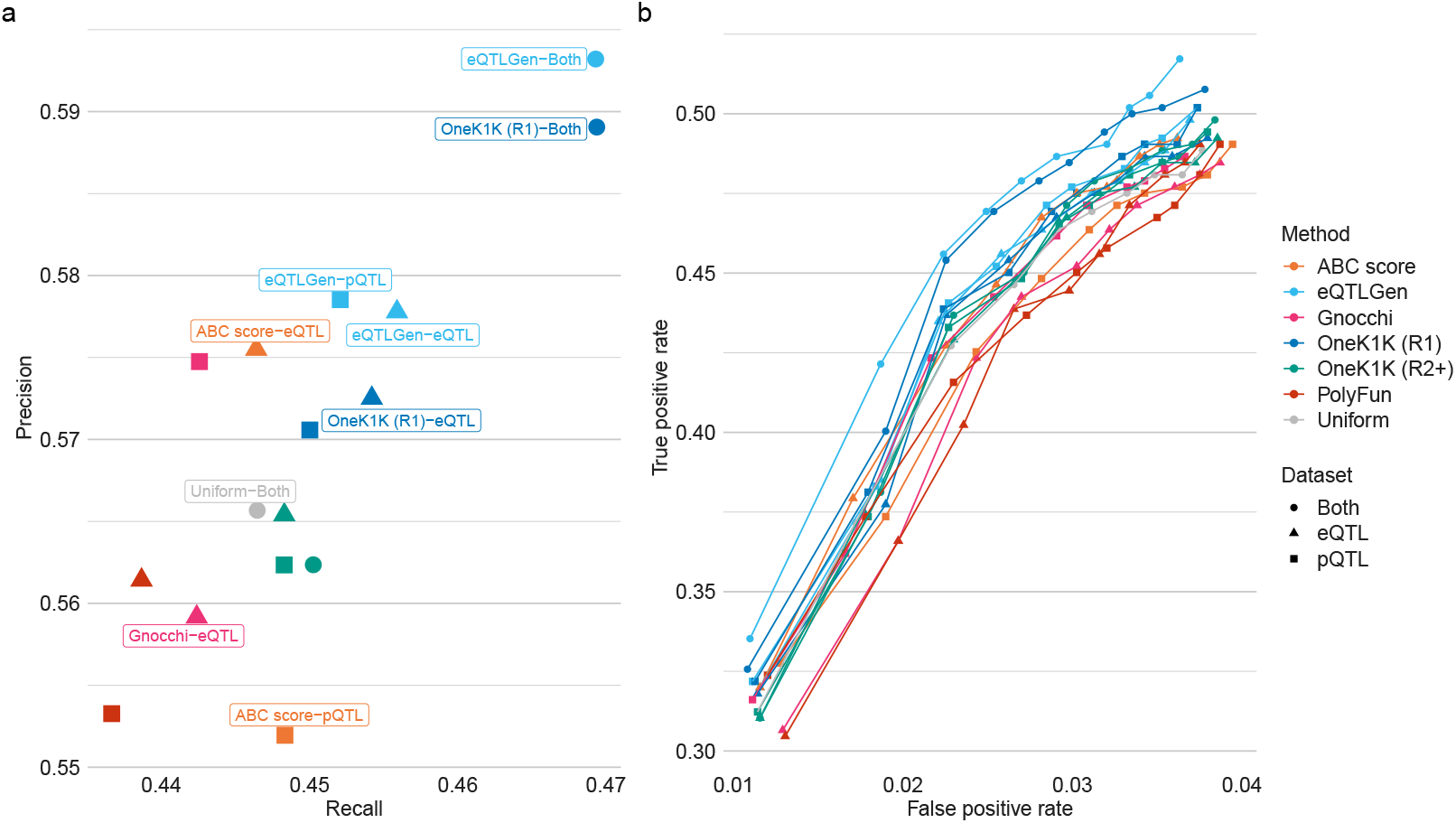
pQTL-eQTL colocalisation performance assessment for coloc with SuSiE. a) Recall and precision of colocalisation at the Pr(*H*_4_) > 0.8 threshold based on ground-truth pQTL-eQTL colocalisations. All 6 different sources of prior information were applied to either the eQTL or pQTL data, with the 3 eQTL-TSS distance density priors additionally applied to both datasets. b) A ROC-style analysis of the ground-truth pQTL-eQTL data using the thresholds Pr(*H*_4_) thresholds: 0.5, 0.55, …, 0.95. In both panels coloc using SuSiE (coloc-susie) is used

**Supplementary Figure S5:**
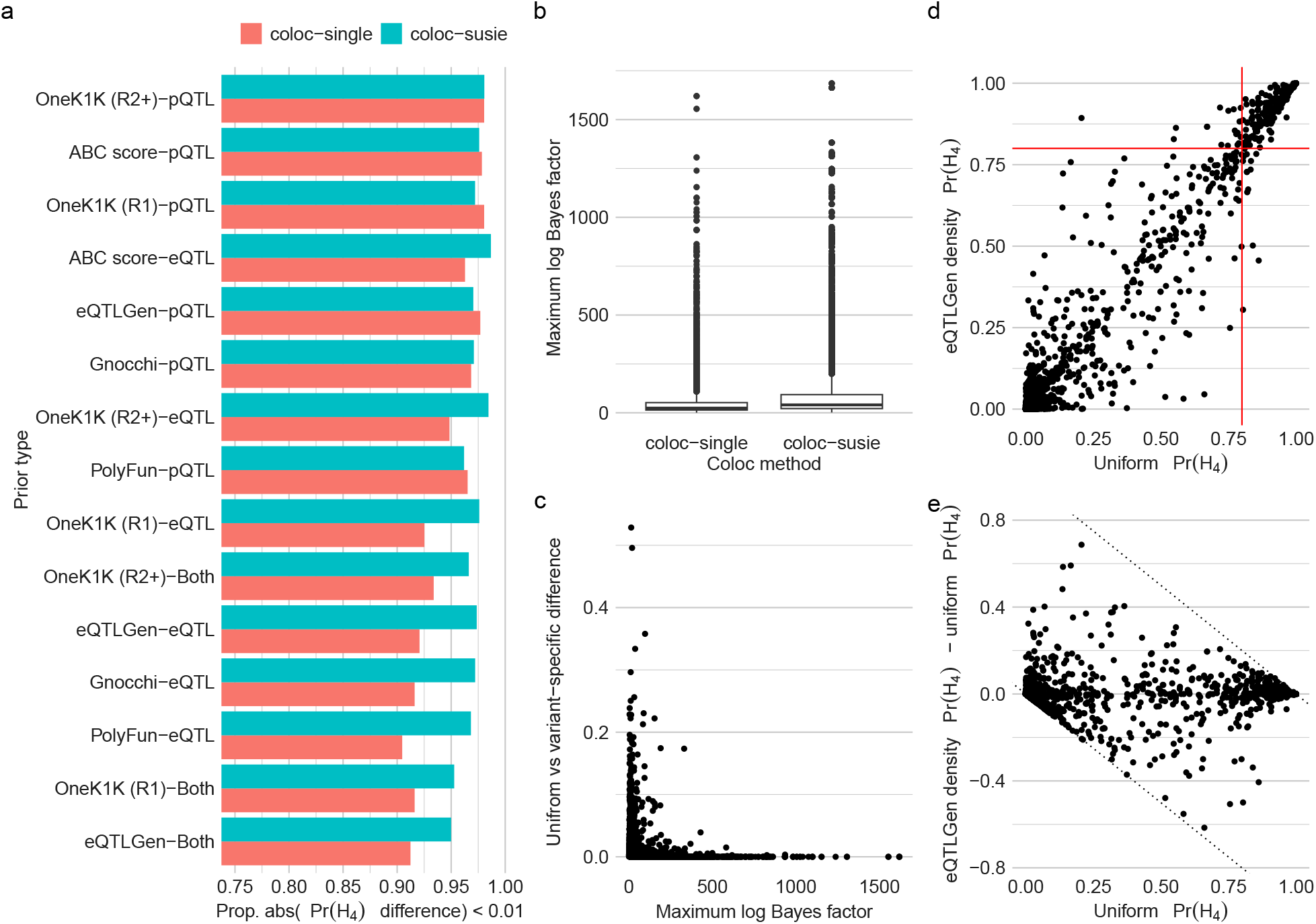
Impact of priors on posterior probabilities in the pQTL-eQTL comparison. a) Plot of the proportion of loci with absolute value change in Pr(*H*_4_) less than 0.01 for coloc method with all priors b) Boxplot of maximum log Bayes factor calculated by coloc-single or calculated by SuSiE and used as input to coloc-susie across loci. c) Scatter plot of maximum log Bayes factor for coloc-single method against the difference in uniform and variant-specific priors, taking the median over prior information sources. Each point is a locus. d) Scatter plot of Pr(*H*_4_) calculated with a uniform prior vs with an eQTLGen-estimated eQTL-TSS density prior across all tested pQTL-eQTL loci for which colocalisation was performed. The red lines show the 0.8 significance threshold for both coloc with uniform and variant-specific priors e) Scatter plot of Pr(*H*_4_) with uniform priors vs the difference between Pr(*H*_4_) with uniform and eQTLGen-estimated eQTL-TSS density prior across all pQTL-eQTL loci where colocalisation was performed. The grey lines show the maximum possible values of the difference in colocalisation probabilities given the value of *Pr*(*H*_4_) calculated with uniform prior probabilities.

