## Supplementary Text for "Variant-specific priors in colocalisation analysis"

#### 1 Summary of coloc method

The coloc methodology assumes that the observed data is generated under one of five hypotheses:

- $H_0$: no association
- $H_1$: association with trait 1 only
- $H_2$: association with trait 2 only
- $H_3$: association with both traits, distinct casual variants
- $H_4$: association with both traits, shared causal variants.

The output of coloc is the posterior probability of these hypotheses for a given region. Interest most often centres on $\Pr(H_4)$ which the probability that the traits do colocalise.

The parameter of the coloc model is a tuple $\boldsymbol{s} = (\boldsymbol{s}^{(1)}, \boldsymbol{s}^{(2)})$ of binary vectors, termed 'configurations', which indicate whether each SNP is associated with each trait. coloc makes the simplifying assumption that that at most one variant is associated with each trait, known as the 'single causal variant' assumption. Therefore, we have $\boldsymbol{s}^{(1)} \in \{\{0,1\}^Q \mid \sum_{i=1}^{Q} s_i^{(1)} \leq 1\}$, and similarly for $\boldsymbol{s}^{(2)}$.

All combinatorially possible 'configurations' can be grouped into five sets $S_0, S_1, S_2, S_3, S_4$ each associated with one of the possible hypotheses. This grouping allows us to compute the probability of each hypothesis by summing the probability of each configuration. Define $\boldsymbol{e}_i$ to be the vector of all zeros with 1 in the $i$th position. Then, these sets are,

- $S_0 = \{\boldsymbol{s}_0\} = \{(\boldsymbol{0}, \boldsymbol{0})\}$
- $S_1 = \{(\boldsymbol{s}, \boldsymbol{0}) \mid \boldsymbol{s} \in \{\boldsymbol{e}_1, \dots e_Q\}\}$
- $S_2 = \{(\boldsymbol{0}, \boldsymbol{s}) \mid \boldsymbol{s} \in \{\boldsymbol{e}_1, \dots e_Q\}\}$
- $S_3 = \{(\boldsymbol{e}_i, \boldsymbol{s}) \mid \boldsymbol{s} \in \{\boldsymbol{e}_1, \dots e_Q\} \setminus \boldsymbol{e_i},\ i \in [1, \dots Q]\}$
- $S_4 = \{(\boldsymbol{e}_i, \boldsymbol{e}_i) \mid i \in [1, \dots, Q]\}$.

To compute the probability of $H_h$, coloc uses the following approach. $\Pr(H_h \mid D)$ is calculated as

$$\begin{aligned}\Pr(H_h \mid D) &= \frac{\Pr(H_h \mid D)}{\Pr(H_0 \mid D) + \Pr(H_1 \mid D) + \Pr(H_2 \mid D) + \Pr(H_3 \mid D) + \Pr(H_4 \mid D)} \\ &= \frac{\frac{\Pr(H_h|D)}{\Pr(H_0|D)}}{1 + \frac{\Pr(H_1|D)}{\Pr(H_0|D)} + \frac{\Pr(H_2|D)}{\Pr(H_0|D)} + \frac{\Pr(H_3|D)}{\Pr(H_0|D)} + \frac{\Pr(H_4|D)}{\Pr(H_0|D)}},\end{aligned}$$

noting in the first line that the denominator sums to 1, and dividing through by $\Pr(H_0 \mid D)$ in the second line. Each fraction is then computed as:

$$\begin{aligned}\frac{\Pr(H_h \mid D)}{\Pr(H_0 \mid D)} &= \frac{\Pr(D \mid H_h)\Pr(H_h)}{\Pr(D \mid H_0)\Pr(H_0)} \\ &= \frac{\sum_{\boldsymbol{s} \in S_h} \Pr(D \mid \boldsymbol{s})\Pr(\boldsymbol{s})}{\Pr(D \mid \boldsymbol{s}_0)\Pr(\boldsymbol{s}_0)} \\ &= \sum_{\boldsymbol{s} \in S_h} \frac{\Pr(D \mid \boldsymbol{s})}{\Pr(D \mid \boldsymbol{s}_0)} \times \frac{\Pr(\boldsymbol{s})}{\Pr(\boldsymbol{s}_0)}\end{aligned}$$

where final expression, the first ratio is a Bayes Factor comparing each configuration to the null configuration ($\boldsymbol{s}_0$) and the second ratio is the prior odds of the configuration compared to the null configuration, $\boldsymbol{s}_0$. To specify the prior odds of each configuration coloc model has three key prior parameters:

- $p_1$: The probability that a variant is causal for trait 1 only;
- $p_2$: The probability that a variant is causal for trait 2 only; and,
- $p_{12}$: The probability that a SNP is causal for both traits.

The prior probability of each configuration is proportional to the product of 1) the prior probability of the causal variant, which may be causal for for either or both traits, and 2) the prior probability of the non-causal variants. These quantities can then be normalised by the their sum across configurations, but as coloc uses the prior odds this step is unnecessary. Precisely, we have that $\Pr(\boldsymbol{s}) \propto$

- $p_0^Q, \ \forall \boldsymbol{s} \in S_0$;
- $p_1^{Q-1} \times p_1, \ \forall \boldsymbol{s} \in S_1$;
- $p_0^{Q-1} \times p_2, \ \forall \boldsymbol{s} \in S_2$;
- $p_0^{Q-2} \times p_1 \times p_2, \ \forall \boldsymbol{s} \in S_3$, and
- $p_0^{Q-1} \times p_{12}, \ \forall \boldsymbol{s} \in S_4$.

Then, the prior odds, $\frac{\Pr(\boldsymbol{s})}{\Pr(\boldsymbol{s}_0)}$, of each configuration in each set relative to the single baseline configuration are

- $1, \ \forall \boldsymbol{s} \in S_0$;
- $\frac{p_0^{Q-1} \times p_1}{p_0^Q} = \frac{p_1}{p_0} \approx p_1 \ \forall \boldsymbol{s} \in S_1$;
- $\frac{p_0^{Q-1} \times p_2}{p_0^Q} = \frac{p_2}{p_0} = p_2, \forall \boldsymbol{s} \in S_2$;

- $\frac{p_0^{Q-2}\times p_1\times p_2}{p_0^Q} = \frac{p_1\times p_2}{p_0^2} \approx p_1 \times p_2,\ \forall \boldsymbol{s} \in S_3$, and
- $\frac{p_0^{Q-1}\times p_{12}}{p_0^Q} = \frac{p_{12}}{p_0} \approx p_{12},\ \forall \boldsymbol{s} \in S_4$,

where we assume that $p_0 \approx 1$ and $p_0^2 \approx 1$. Let $\mathrm{ABF}_i^{(k)}$, denote the approximate Bayes factor for the $i$th SNP, $i \in [1,\ldots,Q]$. in the $k$th trait, $k \in \{1,2\}$. Then, combining the Bayes factors and the prior odds to give $\frac{\Pr(D|H_h)}{\Pr(D|H_0)}$ yields,

- $1,\ \forall \boldsymbol{s} \in S_0$,
- $p_1 \times \sum_{i=1}^{Q} \mathrm{ABF}_i^{(1)}, \forall \boldsymbol{s} \in S_1$
- $p_2 \times \sum_{i=1}^{Q} \mathrm{ABF}_i^{(2)}, \forall \boldsymbol{s} \in S_2$
- $p_1 \times p_2 \times \sum_{\substack{i,j=1 \\ i\neq j}}^{Q} \mathrm{ABF}_i^{(1)}\mathrm{ABF}_j^{(2)},\ \forall \boldsymbol{s} \in S_3$
- $p_{12} \times \sum_{i=1}^{Q} \mathrm{ABF}_i^{(1)} \times \mathrm{ABF}_i^{(2)},\ \forall \boldsymbol{s} \in S_4$.

### 2 Variant-specific prior probabilities calculation

In version 6.0.0 of coloc, the `coloc.abf()` and `coloc.susie()` functions in `coloc` have been updated to include two new arguments: `prior_weights1`, `prior_weights2` which specify non-negative weights for the probability of a variant being causal for trait 1 and trait 2 respectively. These weights are used to calculated variant specific prior probabilities, according to the formula:

$$p_{k,i} = Q \times p_k \times \frac{w_{k,i}}{\sum_{i=1}^{Q} w_{k,i}},\ k \in \{1,2\},\ i \in \{1,\ldots,Q\} \tag{1}$$

where $w_{k,i}$ is the provided weight for SNP $i$ and trait $k$, $Q$ is the number of variants analysed (i.e. represented in the association data for both traits), and $p_{k,i}$ is the prior probability of SNP $i$ and trait $k$. The variant-specific prior probabilities of being causal for each trait are then used to calculate the variant-specific prior probability of causality for both traits:

$$p_{12,i} = \frac{p_{12}}{p_1 p_2} \times p_{1,i} \times p_{2,i} \tag{2}$$

where $p_{12}$, $p_1$ and $p_2$ are as described above.

### 3 Variant-specific prior calculation considerations

#### 3.1 Prior applied to one trait

This calculation of the prior probabilities is designed to ensure that the variant-specific prior probabilities satisfy two conditions: that they are proportional to $w_{k,i}$, and that they encode the same hypothesis prior probabilities as uniform priors. To see that this is the case, note that with uniform priors,

$$\Pr(H_k) \approx Q \times p_k,\ k \in \{1,2\} \tag{3}$$

while, with variant specific priors,

$$\Pr(H_k) = \sum_{i=1}^{Q} p_{k,i} = \sum_{i=1}^{Q} Q \times p_k \times \frac{w_{k,i}}{\sum_{i=1}^{Q} w_{k,i}} = Q \times p_k,\ k \in \{1,2\} \tag{4}$$

Therefore, the variant-specific priors incorporate information from the supplied variant-specific weights while not affecting the overall prior probability of the colocalisation hypotheses. When one of the priors, $p_1$ say, is variant-specific then we have

$$\Pr(H_4) = \sum_{i=1}^{Q} p_{12,i} = \sum_{i=1}^{Q} \frac{p_{12}}{p_1 p_2} \times p_{1,i} \times p_2 = \sum_{i=1}^{Q} \frac{p_{12}}{p_1} \times Q \times p_1 \times \frac{w_{1,i}}{\sum_{i=1}^{Q} w_{1,i}} = Q \times p_{12},$$

so the overall prior weight on $H_4$ is the same as with uniform priors. We describe the case where both priors are variable in methods.

#### 3.2 Prior applied to both traits

In the implementation of variant-specific priors in coloc it is possible to specify variant-specific priors on both traits. Here to simplify notation we let,

$$\pi_{1,i} = \frac{w_{1,i}}{\sum_{i=1}^{Q} w_{1,i}} \text{ and } \pi_{2,i} = \frac{w_{2,i}}{\sum_{i=1}^{Q} w_{2,i}}.$$

In this case, the prior probability of $\Pr(H_4)$ is

$$\begin{aligned}
\Pr(H_4) &= \sum_{i=1}^{Q} p_{12,i} \\
&= \sum_{i=1}^{Q} \frac{p_{12}}{p_1 p_2} \times p_{1,i} \times p_{2,i} \\
&= \sum_{i=1}^{Q} \frac{p_{12}}{p_1 p_2} \times Q \times p_1 \times \pi_{1,i} \times Q \times p_2 \times \pi_{2,i} \\
&= Q \times p_{12} \times \left( Q \sum_{i=1}^{Q} \pi_{1,i} \times \pi_{2,i} \right).
\end{aligned}$$

We consider two scenarios to determine whether this expression is likely to be close to $Q \times p_{12}$. First, if we consider the weights as independent random variables with $\pi_{1,i}$ independent of $\pi_{2,i}$ and $\mathbb{E}(\pi_{1,i}) = \mathbb{E}(\pi_{2,i}) = \frac{1}{Q}$ then we have,

$$\mathbb{E}(\Pr(H_4)) = Q \times p_{12} \times \left( Q \sum_{i=1}^{Q} \mathbb{E}(\pi_{1,i} \times \pi_{2,i}) \right) = Q \times p_{12} \times \left( Q \sum_{i=1}^{Q} \frac{1}{Q^2} \right) = Q \times p_{12},$$

so we recover $Q \times p_{12}$ in expectation. Alternatively, if the weights are exactly equal, so $\pi_{1,i} = \pi_{2,i} = \pi_i$, then

$$Q \sum_{i=1}^{Q} \pi_{1,i} \times \pi_{2,i} = Q \sum_{i=1}^{Q} \pi_i^2 = Q \sum_{i=1}^{Q} (\pi_i - \frac{1}{Q})^2 + 1$$

which is minimised when $\pi_i = \frac{1}{Q}$, $\forall i$, that is when the prior is uniform, in which case it equals 1 so $\Pr(H_4) = Q \times p_{12}$. As the prior deviates from uniformity, the expression increases, indicating increasing the prior weight on $H_4$. This behaviour is sensible as applying the same non-uniform prior to both traits encodes that we believe that their causal variants should lie in a similar location increasing the probability of colocalisation.

### 4 Incorporation of variant-specific prior probabilities into coloc

Prior probabilities are incorporated in the computation of $\frac{\Pr(D|H_h)}{\Pr(D|H_0)}$. Specifically, using the variant-specific prior probabilities we compute $\frac{\Pr(D|H_h)}{\Pr(D|H_0)}$ to be,

- $1,\ \forall \boldsymbol{s} \in S_0$,
- $\sum_{i=1}^{Q} p_{1,i} \times \mathrm{ABF}_i^{(1)}, \forall \boldsymbol{s} \in S_1$
- $\sum_{i=1}^{Q} p_{2,i} \times \mathrm{ABF}_i^{(2)}, \forall \boldsymbol{s} \in S_2$
- $\sum_{\substack{i,j=1 \\ i \neq j}}^{Q} p_{1,i} \times p_{2,i} \times \mathrm{ABF}_i^{(1)} \mathrm{ABF}_j^{(2)},\ \forall \boldsymbol{s} \in S_3$
- $\sum_{i=1}^{Q} p_{12,i} \times \mathrm{ABF}_i^{(1)} \times \mathrm{ABF}_i^{(2)},\ \forall \boldsymbol{s} \in S_4$.
